# Bacteriophage replication strategies are associated with organic matter energy content on coral reefs

**DOI:** 10.1101/2025.05.12.653515

**Authors:** Natascha S Varona, Lisa Schellenberg, Will Barnes, Yun Scholten, Andreas F Haas, Cynthia B Silveira

## Abstract

Bacteriophages, viruses that infect bacteria, play a crucial role in carbon cycling within marine environments. In coral reefs, dissolved organic matter (DOM) released by benthic primary producers such as algae fuels heterotrophic microbial growth, which can be detrimental to corals. While this microbialization process has been associated with the abundance and replication strategies of bacteriophages, the direct relationship between reef DOM composition and bacteriophage communities remains unclear. Here, we combine metabolomics, metagenomes, and viromes to demonstrate that phage abundances have significant relationships with DOM composition on the reefs of Curaçao, Southern Caribbean. By constructing co-occurrence networks between free or cell-associated viruses and exometabolites, we identified thousands of statistically significant associations between phages and organic compounds. While total viral abundances did not significantly correlate with overall dissolved organic carbon (DOC) concentration, cell-associated phages had significantly more positive associations with compounds that had a reduced nominal oxidative state of carbon (NOSC). Furthermore, temperate phages were more frequently correlated with metabolites exhibiting higher Gibbs energy than lytic phages. Six of the ten viruses with the highest number of positive associations with metabolites were temperate (i.e., encoded an integrase or were identified as a prophage), despite this network consisting of approximately 90% lytic viruses. These temperate viruses were predicted to infect members of the genus *Sphingobium*. Together, these findings reveal a connection between phage replication strategies and DOM energy availability with potential implications for coral reef biogeochemistry.

**Significance:** Coral reefs are highly dynamic ecosystems where microbial communities and organic matter cycles are intricately linked. This study provides new insights into how bacteriophages interact with dissolved organic matter (DOM) composition, revealing that cell-associated bacteriophages, particularly temperate phages, are associated with more energy-rich organic compounds. These findings suggest that DOM may have a bottom-up influence on the lysis-lysogeny decision of temperate phages infecting their host or that lysogeny may play an underappreciated role in shaping the reef carbon cycle. Energy-rich organic compounds have generally been associated with increased algal abundances and coral decline. By demonstrating significant connections between viral infection strategies and the energy content of DOM, our results highlight the potential for phages to influence coral reef biogeochemistry and health.

## Introduction

Viruses play crucial roles in marine ecosystems by driving carbon cycling through cell lysis (1–4). It’s estimated that every 0.3 seconds, there are 3 × 10^22^ viral infections occurring in the ocean (2). Bacteriophage-driven lysis of microorganisms releases organic matter and nutrients into the environment, diverting energy from the secondary consumers of the food web (for example, zooplankton), towards predominantly heterotrophic bacteria, a process known as the viral shunt (3). Additionally, lysed organic matter can aggregate with cell debris and mucus, forming marine snow and accelerating carbon export to the ocean floor (5). Viral lysis is estimated to release approximately 145 Gt of carbon per year in the tropical and subtropical oceans (1).

In coral reef ecosystems, carbon fluxes have been shown to influence reef health (6–8). The Disease, Dissolved organic matter, Algae, and Microorganisms (DDAM) model (9, 10) describes a positive feedback loop in which increased dissolved organic matter (DOM) released by fleshy algae fuels heterotrophic microbial growth, leading to oxygen depletion that can cause coral death. The newly open niche space is taken over by algae, which can now perpetuate the DDAM loop by releasing more DOM. Among reef organisms, algae exhibit the highest rates of DOM exudation, with conservative estimates suggesting that turf and macroalgae release approximately 10% of their photosynthetically fixed organic carbon (11, 12). While corals and algae contribute to the majority of reef DOM, algae exude more labile DOM and reduced organic molecules compared to corals (13, 14). These compounds are generally more energy-rich and provide a greater metabolic potential per carbon (15–17), which affects the microbial community composition in reef waters (13, 14, 18, 19). DOM from macroalgae promotes decreased microbial diversity and the growth of copiotrophic taxa such as *Gammaproteobacteria* (20). In contrast, DOM from corals is associated with high microbial diversity, with communities enriched in *Alphaproteobacteria* (20). A study that analyzed over 400 samples from 60 coral reef sites revealed that the microbial consumption of algal-derived DOM leads to an increase in microbial biomass and a shift in community composition, a phenomenon described as coral reef microbialization (21). The algae-derived exudates reduce bacterial growth efficiency to as low as 6%, whereas coral exudates can be metabolized more efficiently at 18% (20). This is because the energy-rich compounds released by algae are not metabolized as efficiently, potentially increasing biological oxygen demand and contributing to microbially driven deoxygenation (11, 22, 23). These combined effects further shift reef community dynamics in favor of fleshy and turf algae (9, 10).

Viruses may be an integral part of the DDAM pathway (24). They may do so by directly lysing different bacterial members of the community (7, 25–27), by releasing DOM into the environment via cell lysis (5, 6), which in turn selects for different microbes, or through different lifecycle strategies (lysis and lysogeny) that may select for a specific community composition. Healthy coral reefs are typically associated with high viral predation pressure (7), with phages frequently targeting *Gammaproteobacteria* (26). However, as bacterial abundance increases, viral replication strategy shifts toward lysogeny, where the viral genome integrates into the host chromosome rather than lysing the cell (25). Through lysogeny, viruses can influence bacterial gene expression, potentially altering microbial functions within the reef ecosystem. For instance, phages in coral reefs have been found to encode genes related to energy, amino acid, and carbohydrate metabolism, as well as the biosynthesis of secondary metabolites, all of which can impact host metabolism (28). Viral reprogramming can modify key metabolic pathways, such as the pentose phosphate pathway, tricarboxylic acid cycle, and glycolysis (29, 30). Additionally, phages can introduce virulence genes (31, 32) and antibiotic resistance genes (33), further influencing and altering microbial dynamics.

While the pairwise relationships between phages and bacteria and between bacteria and DOM have been described in coral reefs, the direct relationship between phages and DOM remains largely unexplored. Understanding this relationship is of particular importance given that DOM sources could have bottom-up influences on viral infection through changes in host metabolism, or that viruses can top-down control DOM release through viral lysis (6) or manipulate host metabolism, affecting DOM composition in the environment.

To investigate the relationship between phages and DOM composition on coral reefs, this study integrates total DOC and Nitrogen concentrations, microscopy-derived cell and viral abundances, size-fractionated metagenomics and viromics, and exometabolomic data from coral reefs in Curaçao. To determine the energy availability of organic compounds, we calculated the nominal oxidation state of carbon (NOSC) per compound, which can be used as a proxy to establish bioenergetic potential for catabolism (34). By constructing a co-occurrence network between organic compounds and phage abundances derived from metagenomes and viromes, we identified thousands of significant associations, revealing a significant relationship between temperate viruses and energy-rich compounds.

## Methods

### Sampling

We collected exometabolomes (n = 20), total DOC (n = 15), total Nitrogen (TN; n = 15), viromes (n = 17), metagenomes (n = 18), and microscopy data (n = 16) across 29 distinct sites on the leeward side of Curaçao in the Caribbean (Fig. 1a). Viromes, metagenomes, and microscopy data were previously published in Varona et al. 2024. Total DOC, Nitrogen, and exometabolomes are described here for the first time. The coral reef benthic boundary layer (< 30 cm above the reef benthos) (35) was sampled between 9 and 10 AM via SCUBA at between 5 and 10 m depth in July 2021, as previously described in Varona et al. 2024 (26). Briefly, water samples were collected using bilge pumps attached to 18 L collapsible carboys for metagenomes, microscopy, and viromes. Custom-made Niskin bottles (2L) were used to collect samples for DOC, Nitrogen, and exometabolomes. All samples were transported to CARMABI Research Station and processed within 2 hours of collection. Seven sites have all data types (metagenome, virome, microscopy, DOC, Nitrogen, and exometabolome) while the other sites have gaps in the data availability (Table S1).

**Figure 1.**
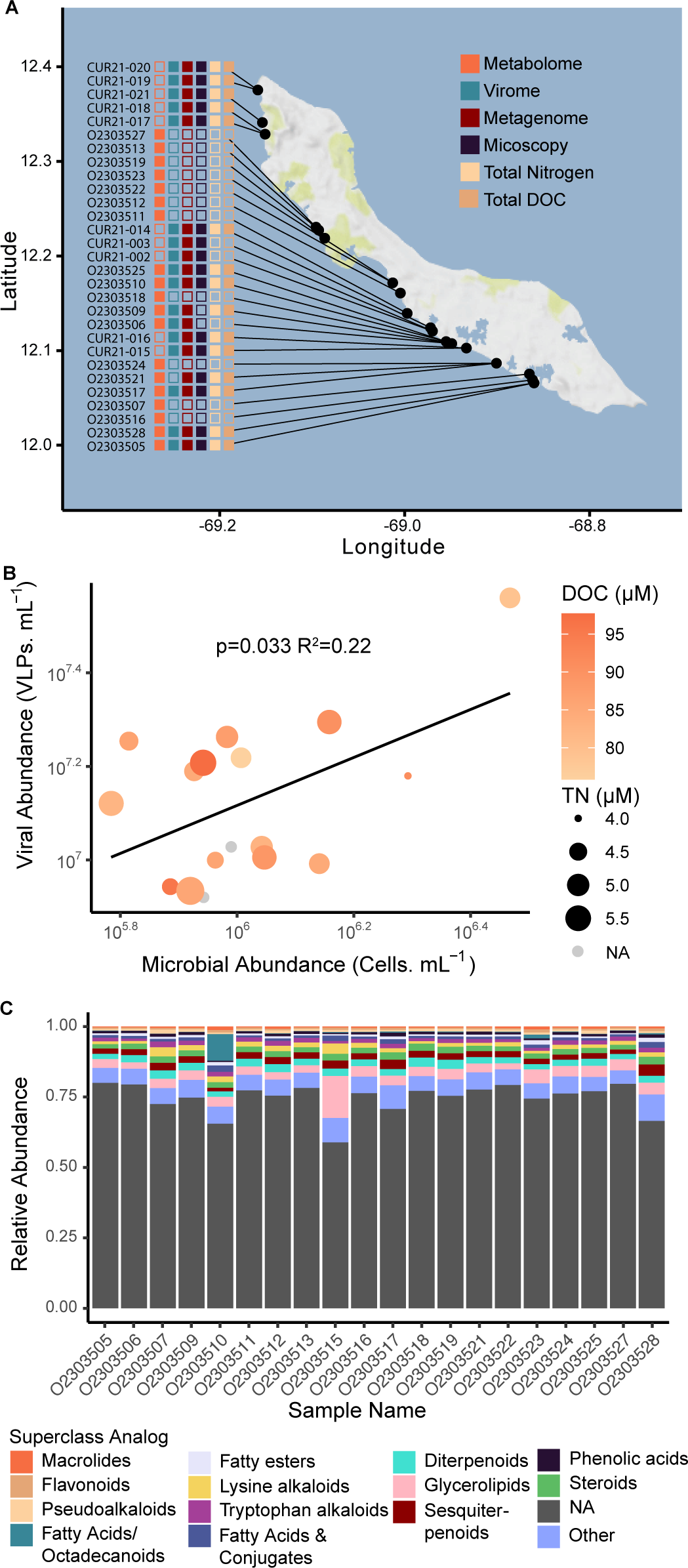
Relationships between viral and microbial abundance and organic matter content. (A) Map of Curaçao showing the 29 sampling sites where exometabolomes, viromes, metagenomes, microscopy, total nitrogen, and total DOC were collected. (B) Linear regression between viral-like-particle abundances and cell abundances, showing a significant positive relationship based on log_10_-normalized data, p-value=0.033, R^2^=0.22. The color intensity indicates DOC concentration, and the size represents the total Nitrogen concentration, which showed no significant correlations with viral-like-particle abundances, cell abundances, or virus-to-microbe ratios. (C) Relative abundance of superclass analogs in the exometabolomes by site.

Upon arrival in the laboratory, 2L seawater subsamples from the carboys were pre-filtered with 8 µm filters (Whatman, Milwaukee, USA) at an approximate flow rate of 12 mL/min using a peristaltic pump Masterflex L/S (Avantor, Gelsenkirchen, Germany), and cells were collected on 0.22 µm Sterivex filters (Millipore Sigma, Burlington, USA) for metagenome sequencing. These filters were stored at −80 °C and kept frozen during transport until DNA extraction with Nucleospin Tissue Kit (Macherey-Nagel, Duren, Germany). For viromes, 5 L subsamples were pre-filtered using 8 µm and 0.45 µm filters and concentrated using a Tangential Flow Filtration system, Vivaflow 100kDa cassette (Sartorius, Goettingen, Germany) (36). Samples were treated with chloroform (0.1%) and stored at 4 °C until precipitation with Polyethylene Glycol 8000 (10%) and DNA extraction with the Purelink^TM^ Viral DNA kit (ThermoFisher, Carlsbad, CA, USA).

### Genome analyses

Sequencing of metagenomes and viromes, viral identification, and abundance calculations were described in Varona et al. 2024 (26). Briefly, total DNA for metagenomes and viromes was sequenced on an Illumina HiSeq 4000 (2×150 bp). Reads were trimmed and quality filtered with BBDuk v.38 (37), assembled into contigs using SPAdes v3.15.5 (38). Six selected samples collected on a 0.22 µm filter underwent MetaHi-C processing with Phase Genomics’ ProxiMeta Hi-C kit (Seattle, WA, USA). Viral genomes were identified using VIBRANT v1.2.1 (39), dereplicated with CD-HIT v4.8.1 (40), binned with vRhyme v1.1.0 (41), and required a minimum length of 10,000 kb or, if smaller, were assessed for high or complete quality with CheckV v1.0.1 (42). Contigs containing contamination, as determined by CheckV, were removed for gene analyses. vMAGs were clustered based on gene content and nucleotide identity using Virathon v.1 (43), and viral abundances were estimated using genome length-adjusted read counts. The lengths and abundances of viral genomes and gene fragments are publicly available through Figshare (44). Viral host predictions for the viruses with the most connections were made using IPhop v1.3.3-0 with the most recent available database (August 2023) and only reported matches with IPhop scores greater than 0.9 (45). Viral lifestyles were established through VIBRANT v1.2.1, which uses criteria for identifying integrases or if viral sequences were determined to be host-integrated (Table S2).

### Dissolved organic matter processing and extraction

The water samples for exometabolomics were processed using solid-phase extraction of dissolved organic matter (SPE-DOM; (46)) with slight modifications (47). Each sample was filtered through a 0.22 μm Sterivex-GP filter (Millipore, Burlington, Massachusetts, USA) at ∼1 L/hour. After discarding the initial 200 mL, 1.1 L of filtrate was collected, acidified to pH 2 with 36% ultrapure HCl, and extracted using a 200 mg Bond Elut-PPL resin cartridge (3 mL; Agilent 12105005). Before the field campaign, PPL cartridges were preconditioned with 3 mL of LC-MS grade MeOH (>=99.9% MeOH; Honeywell, Riedel-de Haën), soaked overnight in MeOH, rinsed twice with de-ionized water, and dried after a final MeOH rinse. Immediately prior to sample processing, PPL cartridges were activated with 3 mL of MeOH and two fills of pH 2 water.

Samples were extracted at ∼5 mL/min using a Masterflex L/S (Avantor, Gelsenkirchen, Germany), and acid-leached tubing (0.1M HCl), as well as flushed with 100–200 mL of sample. Cartridges were desalinated with three fills of pH 2 water and dried with Nitrogen gas. The cartridges were stored at −80°C. Prior to LC-MS/MS analysis, dissolved organic matter (DOM) was eluted with 2 mL LC-MS grade MeOH into combusted 1.5 mL LC glass vials (BGB Analytik AG, Boeckten, Switzerland). The extracts were dried overnight in a vacuum centrifuge, re-dissolved in 100 μL LC-MS grade MeOH, and transferred to a combusted glass insert.

### Exometabolomic analysis

Five µL of DOM extract were injected into an ultra-high performance liquid chromatography system (UHPLC) (Agilent 1290 Infinity II), equipped with a thermostatted auto-injector and column compartment, coupled to a Q-Exactive Plus Orbitrap Mass Spectrometer (Thermo Fisher Scientific, Bremen, Germany) equipped with an Ion-max source with a HESI (Heated Electrospray Ionization) probe according to the protocol of Petras et al., 2017 (48). Chromatographic separation was performed using a C18 core-shell column (Kinetex, 150 × 2 mm, 1.8 µm, 100 Å; Phenomenex) at 30°C with a flow rate of 0.5 mL/min. Solvent A: H2O + 0.1% formic acid (FA); Solvent B: acetonitrile + 0.1% FA. Elution program: 0.5 min at 5% B, linear gradient to 50% B over 7.5 min, to 99% B over 2 min, 7 min wash at 99% B, and 8 min re-equilibration. Ion source settings were as following: sheath gas 70 AU, auxiliary gas 15 AU, probe temperature 400°C, spray voltage 3.5 kV, capillary temperature 320°C, S-Lens 50V. The maximum ion injection time was 100 ms with automated gain control (AGC) targets of 1×10^6^ for MS1 and 3.0×10^5^ for MS/MS, with a 10% minimum C-trap AGC threshold. Compounds were detected in the m/z 150–1500 range at 70,000 resolution (m/z 200) and one micro-scan per MS1. MS/MS spectra were acquired in DDA mode for the top 5 abundant ions, with 17,500 resolution and one micro-scan. Fragmentation used a stepwise collision energy (20-40%) with z = 1. MS/MS experiments were triggered 2-15 sec after peak apex, with dynamic exclusion set to 5 sec and 3 ppm mass tolerance. Quality controls included a Sulfa mix (0.01 µg/mL; Agilent STD6.1, sulfamethizol, sulfamethazine, sulfachloropyridazine, sulfadimethoxine), and a control seawater sample at the start and end of the sequence as well as MeOH injections at the start, end, and between every 5 samples.

The data from liquid chromatography-tandem mass spectrometry were processed in MZmine3 (49), followed by spectral matching with GNPS (https://gnps.ucsd.edu) (50) using the feature-based molecular networking workflow (FBMN) (51). MS/MS raw datafiles were converted to .mzML in centroid mode using MSConvert (Proteowizard) and consequenctly processed with MZmine3 (version 3.9.0) (49). Mass detection was performed in centroid mode with a minimum signal-height threshold of 1.0×10^5^ for MS^1^ and 1.0×10^3^ for MS^2^ levels. Features were generated using the ADAP chromatogram builder (52) based on extracted ion chromatograms (XICs) from each detected m/z value, requiring a minimum of 4 consecutive scans. Parameters included a group intensity threshold of 1.0×10^5^, minimum highest intensity of 2.0×10^5^, and scan-to-scan m/z tolerance of 0.0015 m/z or 10 ppm. The local minimum feature resolver was used to separate overlapping or co-eluting peaks.MS/MS scan pairing was activated, the precursor tolerance was set to 0.010 m/z or 10 ppm, and the retention time tolerance was 0.050 min. Chromatographic settings included an 85% threshold, 0.08 min minimum search range, and minimum absolute peak height of 2.0×10^5^. Additional settings included a minimum peak top-to-edge ratio of 1.4, a peak duration range of 0.00–2.0 min, and a minimum of 4 scans. Isotope peak grouping applied a 13C filter with m/z tolerance of 0.001 or 5 ppm, retention time tolerance of 0.1 min, and a maximum charge of 2, selecting the most intense peak as the monoisotopic ion. XIC alignment across samples was performed using the Join aligner with thresholds of 0.0015 m/z or 10 ppm (weight = 3), retention time tolerance of 0.150 min (weight = 1), and mobility weight of 1. Features with at least two isotope peaks, occurring in at least two samples with MS^2^ scans, were retained. Further filtering utilized the multithreaded peak finder with a 10% intensity tolerance, 0.0015 m/z or 10 ppm sample-to-sample tolerance, 0.15 min retention time tolerance, and a minimum of 3 scans. Duplicate peaks were filtered with m/z tolerance of 0.001 or 5 ppm and RT tolerance of 0.05 min. Correlation grouping (metaCorrelate) was applied with a 0.5 min RT tolerance, 3.0E+04 intensity threshold, ≥60% intensity overlap, exclusion of gap-filled features, and feature shape correlation with a minimum of 5 data points and minimum of 2 data points on edge using Pearson’s coefficient with a minimum feature shape correlation of 85%. Ion identity networking was performed with m/z tolerance of 0.0015 m/z or 5 ppm, maximum charge of 2, and maximum cluster size of 2, considering adducts [M-H]⁻, [M+H]⁺, [M+Na]⁺, [M+NH₄]⁺ and modification [M-H₂O]. Refinement steps included removing networks without monomers and applying adducts and modifications: [M+H]+;[M+Na]+;[M+K]+;[M+NH4]+; [M+2H]2+; {M+H+NA]2+; [M+H+NH4]2+; [M-H+2NA]+; [M+Ca-H]+; [M+Fe-H]+ and : [M-H2O];[M-2H2O];[M-3H2O];[M-4H2O];[M-5H2O]. Networks smaller than size 2 were removed, and a link threshold of 4 was applied. Lastly, consensus MS/MS of each feature were exported as .mgf files for molecular networking through GNPS.

A feature-based molecular network was created using workflow version 28.2 on the GNPS platform. The spectral network was built with a 0.01 Da precursor and fragment ion mass tolerance, a minimum cosine score of 0.7, and more than 4 matched peaks. Only the top 15 edges per node were kept, with a maximum connected component size of 200 and a 500 Da precursor shift limit. Consensus spectra were searched against the GNPS library and in analog mode with variable dereplication (50), using a 200 Da maximum mass difference and reporting the top result.

Data clean-up and normalization were performed using a statistical pipeline for Feature-based molecular networks from non-targeted metabolomics data (53). Blanks were removed using a 0.3 threshold, allowing up to 30% blank contribution. Missing values were imputed by replacing zeros with random values between 0 and the minimum value in the blank-removed feature table, based on relative intensity frequencies. Sample-centric normalization (Total Ion Current, TIC) scaled feature intensities by the total ion current of each sample.

The freely available software R version 4.4.2 (2024-10-31 ucrt) in platform: x86_64-pc-linux-gnu running under: Ubuntu 22.04.3 LTS. Installed packages are textclean, rmarkdown, knitr, kableExtra, tictoc, expss, vegan, stringi, psych, nortest, binom, epitools, car, ape, wesanderson, RColorBrewer, data.table, DescTools, broom, readxl, multcomp, scales, reshape, reshape2, cluster, ggfortify, rfPermute, plyr, tidyverse, tibble, dplyr, svglite, dunn.test, UpSetR, gridExtra, grid, ggpubr, rstatix, ComplexUpset, cowplot, scatterplot3d, pdftools, png, magick, devtools, ggplot2. Analysis was done on 25 runs of which 20 were samples and 5 blanks. Contaminants were flagged and removed if *max*(*blanks*)>=*mean*(*peakarea*)∗0.5max(blanks)>=mean(peakarea)∗0.5, thus over all samples. By comparing the datasets before and after gap filing, the peak area background noise level was set on 3595. Transient features are features that do not pass the background noise in at least 2 samples. These transient features were removed. Feature peak areas were normalized by *log*10(*peakarea*)log10(peakarea) transformation. For all statistical analysis on normalized peak areas, a dataset was used where all peak areas that were replaced by 1000 + random number between 0 and 1 before the transformation. GNPS annotated 2703 features as a known compound based on their MS^2^ spectra. Another 14078 features were matched to highly similar MS^2^ spectra, so called Analog hits. These analog hits might provide structural information on the unknown compounds. The nominal oxidation state of carbon (NOSC) values were calculated based on the predicted formulas, which were derived from the compounds’ Simplified Molecular Input Line Entry System (SMILES) annotation and converted to a chemical formula using the R package cdk. NOSC and Gibbs Energy were calculated based on the number of the elements C, H, N, O, P, and S, following previously published equations (15, 34). Exometabolite abundances, GNPS matches, analog predictions, and NOSC values are publicly available through Figshare (54).

### Network analyses

To identify associations between viruses and compounds, Pearson correlation networks were constructed for virome (n=7, Table S3) and metagenome (n=8, Table S4) datasets separately. Only viruses that were present in at least five sites were included in the analysis. Pearson’s values were corrected using the Benjamini-Hochman method for multiple comparisons, and only associations with corrected p-values < 0.01 and correlation coefficients greater than an absolute value of 0.95 were kept for network analysis. Network statistics were generated through Cytoscape v.3.10.3 *Analyze network* function. In downstream analyses, to group compounds by class, the compound class was prioritized; if the compound class was unknown, the analog class was used. All downstream statistical analyses were performed in RStudio Version 2022.12.0+353. For association testing, Chi-squared tests were applied to the relationships between viral lifestyle and compound class. These tests were performed using Monte Carlo simulations, with 50,000 replicates.

### Data Availability

Raw sequence reads are available through the NCBI Sequence Read Archive (SRA) under PRJNA975592. Virus and exametabolome data are available through Figshare as follows: fasta file with viral genome sequences (vMAGs and contigs) https://doi.org/10.6084/m9.figshare.23313773.v1; viral sequence unique ID, length, and abundance https://doi.org/10.6084/m9.figshare.28924508.v2; exometabolite unique ID and abundance https://doi.org/10.6084/m9.figshare.28924043.v2. Bash scripts used to process sequence data are available via GitHub (https://github.com/Silveira-Lab/Varona-et-al-2023-Phage-host-network-structures).

## Results

### Exometabolomic, viral, and bacterial profiles of coral reefs

DOC concentrations ranged from 75.8 µM to 86.3 µM, while total Nitrogen (TN) ranged from 4.0 µM to 5.9 µM (Fig. 1b). Viral-like-particle abundance derived from microscopy exhibited a significant linear relationship with cell abundance (Linear regression, p-value = 0.033, R² = 0.22) (Fig. 1b). However, total DOC did not exhibit significant correlations with viral-like-particle abundance (Linear regression, p-value = 0.33), cell abundance (Linear regression, p-value = 0.53), or the virus-to-microbe ratio (Linear regression, p-value = 0.77). Similarly, no significant correlations were observed between TN and viral-like-particle abundance (Linear regression, p-value = 0.79), cell abundance (Linear regression, p-value = 0.34), or the virus-to-microbe ratio (Linear Regression, p-value = 0.42). Likewise, the total fractional abundance of viruses in metagenomes and viromes displayed no significant relationship with DOC (Linear regression, p-value = 0.87 for metagenomes; Linear regression, p-value = 0.47 for viromes).

Untargeted metabolomics using Liquid chromatography coupled with tandem mass spectrometry (LC-MS/MS) identified 32,854 unique features (Table S4). GNPS annotations identified 2703 features as known compounds based on their MS² spectra, with an additional 14078 features classified as analog hits (highly similar MS² spectra). These analog hits accounted for approximately 25.3% of all features (Fig. 1c, Table S5). Using the Deep Neural Network-Based Structural Classification Tool, NP Classifier (55), the superclass-level classification of analog hits revealed that glycerolipids constituted the largest fraction (mean = 3.96%, SD = 2.66%), followed by diterpenoids (mean = 2.51%, SD = 0.42%), sesquiterpenoids (mean = 2.28%, SD = 0.60%), and steroids (mean = 2.00%, SD = 0.33%).

### Viruses associate with energy-rich metabolites

To investigate the correlations between viral abundances (from metagenomes and viromes) and metabolite abundances, Pearson correlations yielded 18,619 significant links between individual metabolites and metagenome-derived viruses (hereafter referred to as cell-associated viruses) and 15,199 significant links between individual metabolites and virome-derived viruses (hereafter referred to as free viruses). Most correlations between metabolites and viruses were positive (Fig. 2a). In the cell-associated virus-metabolite network, 4,594 unique metabolites correlated positively, while only 447 were negatively correlated. Similarly, 4,680 unique metabolites were positively associated with free viruses, whereas 554 were negatively associated. Of the positively associated compounds, 2,229 metabolites were shared between metagenome-associated viruses (48.4%) and virome-associated viruses (47.5%). Of the negatively associated compounds, only five metabolites were shared both free and cell-associated viruses.

**Figure 2.**
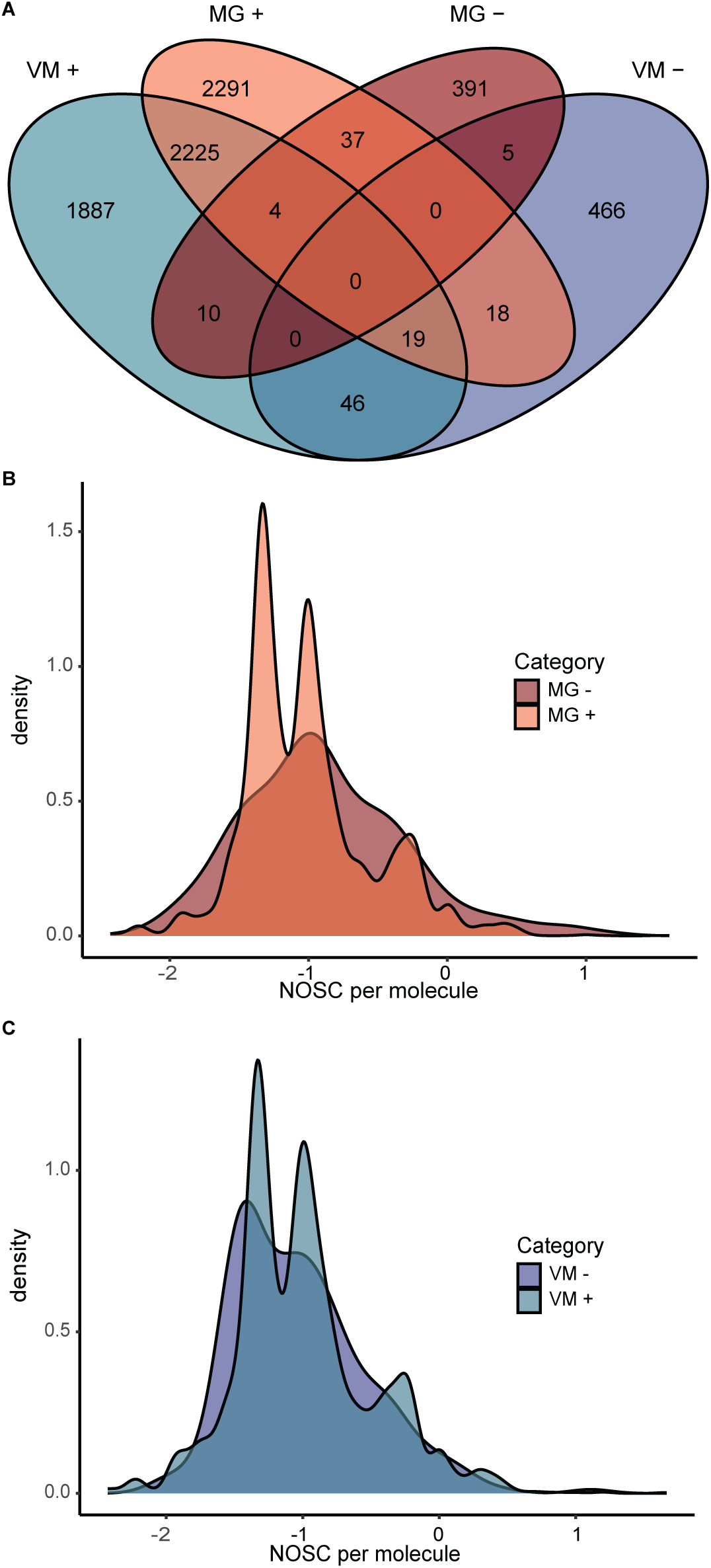
Relationships between viral abundances and metabolites. (A) Venn diagram showing the unique metabolites significantly associated (BH-corrected Pearson’s correlation, p < 0.01) either positively (+) or negatively (−) with viral abundances from metagenomes (MG) and viromes (VM). (B) Density plot comparing the nominal oxidative state of carbon (NOSC) values for metabolites associated with metagenome-derived viral abundances. Positively associated metabolites exhibited significantly more negative NOSC values (mean = −1.02) compared to negatively associated metabolites (mean = −0.888) (Wilcoxon rank-sum test, p-value = 0.000990). (C) Density plot comparing NOSC values for metabolites associated with virome-derived viral abundances. No significant difference in NOSC values was observed between positively and negatively associated metabolites (Wilcoxon rank-sum test, p = 0.140).

For metabolites with feature information annotated via GNPS or analog prediction, we calculated the NOSC as a proxy for the potential energy available during catabolism (15). Cell-associated viruses were more frequently linked to more reduced metabolites (mean NOSC = −1.02) compared to negatively associated metabolites (mean NOSC = −0.888, Wilcoxon rank-sum test, p-value = 0.000990) (Fig. 2b). We identified an increase in density of positive correlations from NOSC values −1.45 to −1.18 with mainly fatty acyls (71.1%, Fig. S1). Of these fatty acyls, 78.4% were linoleic acids and derivatives. A second peak, at NOSC values of approximately −1.10 and −0.88, was comprised of associations with organooxygen compounds (26.6%), prenol lipids (25.0%), steroids and steroid derivatives (15.4%) and fatty acyls (11.1%) (Fig. S2). In contrast to the cell-associated virus-metabolite network, no significant differences in NOSC values were observed in the free viral network (Wilcoxon rank-sum test, p-value = 0.140) (Fig. 2c).

### Correlations between metagenome-derived viral abundances and metabolites

Of the 18,619 significant links in the network connecting metabolites and the metagenome viruses (> 0.45 µm fraction), 18,109 were positive associations (Fig. 3a), and 831 were negatively associated (Fig. S3). The positive network consisted of 5614 nodes, of which 1020 were viruses, and 4594 were metabolites, with an average number of neighbors of 14.75, indicating that viruses are more frequently associated with multiple metabolites with moderate connectivity. The network was fragmented into 635 connected components, meaning it consisted of multiple sub-networks rather than a single, fully interconnected system. This structural organization suggests that certain groups of viruses and metabolites interact independently from others, though this may also result from stringent cut-offs, which would increase false negatives. The network also expresses high heterogeneity, at 6.77, meaning some nodes have significantly more connections than others. To pinpoint which nodes may be keystone features of this network, we further investigated which individual metabolites have the highest degree (most connections to other viruses, Fig. 3b). The 10 most connected metabolites formed 11-15 positive correlations. Although the majority of compounds are unknown, the metabolite with the most positive links was an analog of an organonitrogen compound. Two others could be identified as fatty acyls. One of the fatty acyls (X8968) had a spectral match to 9,12-Octadecadiynoic Acid from NIST14. For metabolites with the highest number of negative links, analog hits to prenol lipids made up three of the six identified compounds, which include organonitrogen compounds, phenols, and steroids and derivatives.

**Figure 3.**
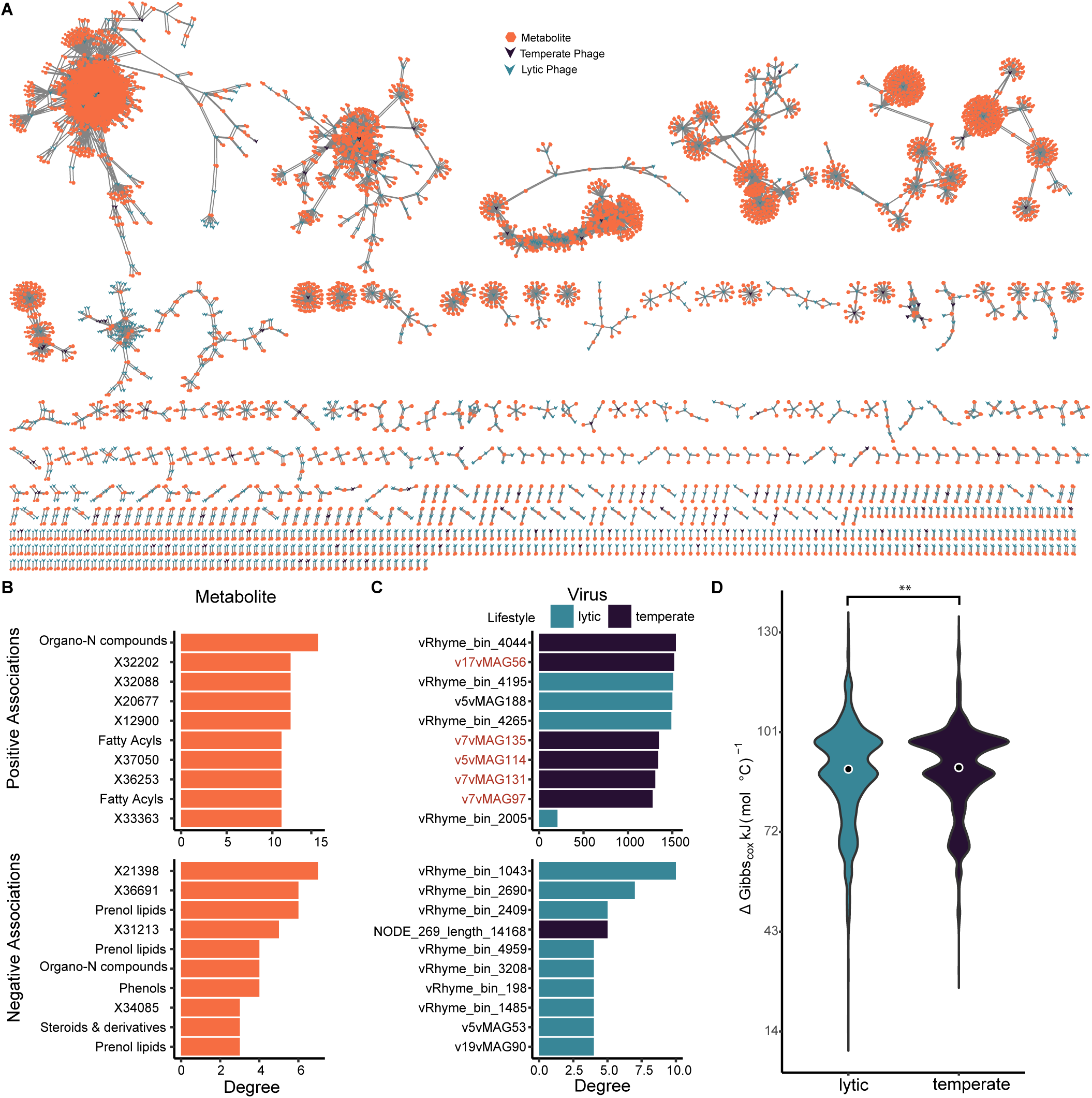
Network of interactions between metagenome-derived phage abundance and metabolites. (A) Network of positive associations between metabolites and phage abundances derived from metagenomes (> 0.22 μm fraction). V-shaped nodes represent phages and hexagon-shaped nodes represent metabolites. Edges represent significant positive correlations (Benjamini-Hochman corrected Pearsons, p-value < 0.01). (B) Metabolites with the most degree (connections) with positive associations (top) and negative associations (bottom). If the compound was a previously known compound or matched an analog, the class was included as the y-axis label. (C) Viruses with the most degrees (connections) with positive associations (top) and negative associations (bottom). Viruses highlighted in red have previously been reported to be associated with the host *Sphingobium yanoikuyae.* (D) Gibbs free energy of metabolites associated with lytic and temperate viruses (Wilcoxon rank-sum test, W=4993937, p-value = 0.00404, lytic mean = 88.8 KJ (mol °C) ^-1^; temperate mean = 89.9 KJ (mol °C) ^-1^).

We also investigated the viruses with the most degrees, which were higher in the positively associated networks than negative networks (Fig. 3b). Though the difference in maximum number of links is influenced by the difference in network size, the negative network was approximately 21 times smaller, while the maximum degrees were 150 times smaller. The viruses with the most positive associations form links with 1541 metabolites. Five of the viruses with the highest number of associations (v17VMAG56, degree = 1521; v7vMAG135, degree = 1349; v7vMAG131, degree = 1309, v7vMAG97, degree = 1278, v5vMAG114, degree = 1341) have been previously found to associate with the host *Sphingobium yanoikuyae* via proximity-ligation (26). Additionally, genome-based predictions of the host of the viruses revealed further potential host taxa (Table S6): vRhyme_4044 was also predicted to infect the host of the genus *Sphingobium,* and three viruses predicted to infect hosts of the class *Bacteroidia* (v5vMAG188), *Coriobacteriia* (vRhyme_4265), and *Alphaproteobacteria (*vRhyme_2005). Three hosts were predicted for negatively associated viruses: *Bacteroidia* (vRhyme_bin_3208, v5vMAG43), *Acidimicrobia* (v19vMAG90), and *Gammaproteobacteria* (vRhyme_bin_1485).

Six of the 10 viruses with the most associations were temperate (i.e., encoding an integrase or identified as a prophage), despite the positive network consisting of 90% lytic viruses. To further investigate differences in the links formed between lytic and temperate viruses, we compared the differences of Gibbs free energy from metabolites correlated with each virus (Fig. 3d). A Wilcoxon-rank sum test revealed a small but significant difference in the distribution of Gibbs energy of compounds associated with lytic versus temperate viruses (Wilcoxon rank-sum test, p-value = 0.00404; lytic mean = 88.8 KJ (mol °C) ^-1^; temperate mean = 89.9 KJ (mol °C) ^-1^). This indicates that temperate viruses tend to associate with energy-rich compounds more so than lytic viruses within the metagenome-virus-metabolite network. To assess whether this pattern held across the core range of the data, we subsampled metabolites with Gibbs energy values between 85.1 and 101.3 kJ mol⁻¹ °C⁻¹, capturing the central mode of the distribution. Within this subset, the Wilcoxon rank-sum test did not detect a statistically significant difference between the two groups (p = 0.0524). However, a Kolmogorov–Smirnov test revealed a significant difference in the overall shape of the distributions (D = 0.061442, p = 0.001347), indicating that the frequency and/or spread of metabolite associations differ between temperate and lytic viruses, with a potential enrichment of temperate virus associations in specific energy ranges.

### Correlations between virome-derived viral abundances and metabolites

A second network was built with viruses from the free virus fraction (virome, < 0.45 µm) and metabolites. The positive virome network had 4972 nodes, of which 4919 were metabolites and 781 were viruses, forming 14,466 positive associations (Fig. 4a), about 1.25 times smaller than the metagenome-derived (>0.22 um) network. Similarly, the negatively associated virome network was smaller than the positive network, with 1059 total nodes and 733 links (Fig. S4). This network also showed a high heterogeneity of 8.71. The most abundant viruses formed a high number of linkages, 110 to 1477, and the metabolites formed 45-59 linkages (Fig. 4b). Three of these viruses encoded metabolic genes: vRhyme_bin_3950, v5vMAG188, and NODE_1_length_62316_cov_79.381459, which were all related to pyrimidine metabolism (deoxyribonucleotide biosynthesis). While the hosts of these viruses have not yet been described, we used iPHop to predict potential hosts, which identified two hosts belonging to the class *Bacterioidia* (vRhyme_bin_4225 and v5vMAG188), one associated with *Bacilli* (vRhyme_bin_4224), and *Gammaproteobacteria* (vRhyme_bin_4584). Of the viruses with the most negative associations, four were predicted to infect bacteria of the class *Bacteroidia,* and two were predicted to infect *Alphaproteobacteria* (Fig. S5a). The metabolic compounds were predominantly unknown, except for one belonging to the analog class benzene and derivatives (Fig. 4 B). This one had a spectral match with fexofenadine (Massbank: EX301902). The metabolites with the most negative associations (Fig. S5b) had an analog hit to steroids and derivatives (X11307), and two known compounds, diethylene glycol monobutyl (X332, GNPS Spectrum ID: CCMSLIB00005726830) and DEET (X1847, GNPS Spectrum ID: CCMSLIB00000579756). This network also differed from the metagenome-derived network, where there were no temperate viruses with a high number of linkages or a significant difference in the Gibbs free energy of compounds correlated with temperate or lytic viruses (Wilcoxon-Rank Sum Test, W = 869807, p-value = 0.857).

**Figure 4.**
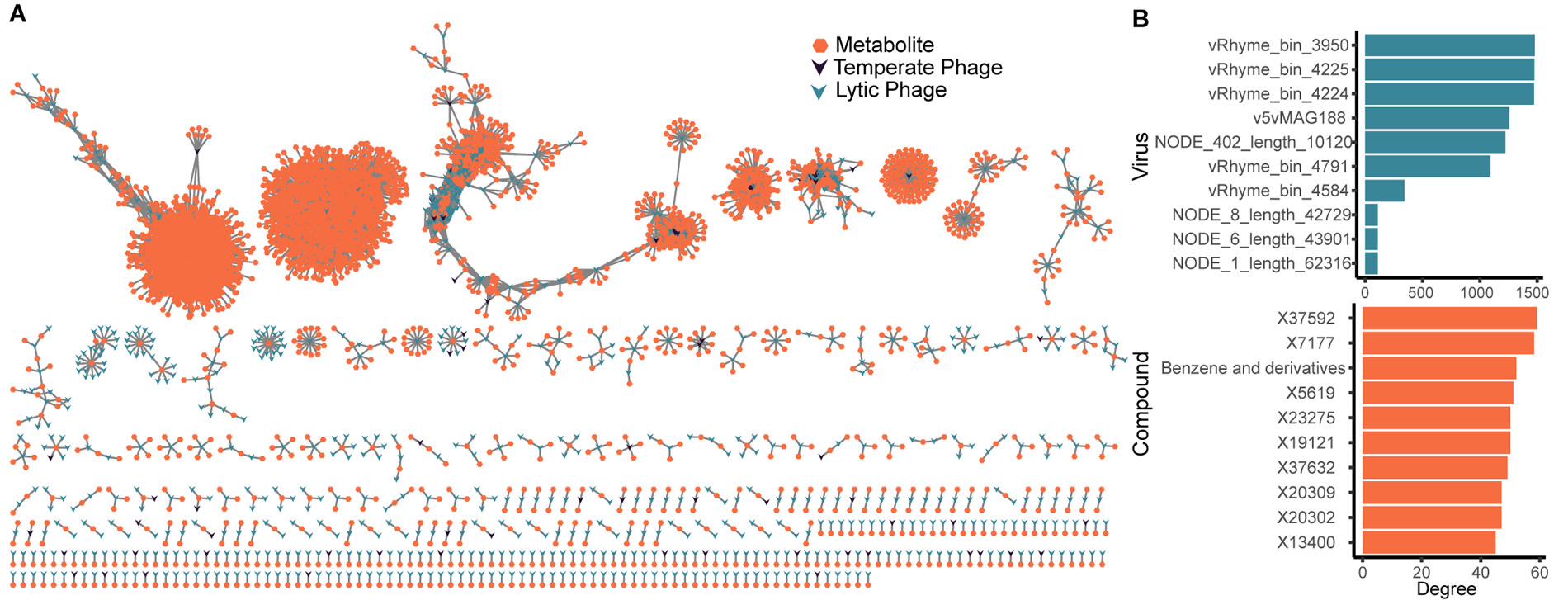
Network of positive associations between free viral abundances and metabolites. (A) Network of positive associations between phages (< 0.45 μm fraction, virome) and metabolites. V-shaped nodes represent phages and hexagon-shaped nodes represent metabolites. Edges represent significant positive correlations (Benjamini-Hochman corrected Pearsons, p<0.01). (B) Viruses with the most degrees (connections) with positive associations (top) and metabolites with the most degree (connections) with positive associations (bottom).

### Differences between compounds associated with temperate and lytic viruses

To further investigate differences between compounds associated with lytic and temperate viruses, we performed a Chi-squared test with Monte Carlo simulations (50,000 repetitions). The analysis revealed a significant association between viral lifestyle (lytic vs. temperate) and the compounds they interact with in both the positive metagenome-associated viruses-metabolite network (χ² = 312.13, p-value < 0.0001) and the virome-associated viruses-metabolite network (χ² = 588.27, p-value < 0.0001). To identify the specific groups of compounds driving these differences, we examined standardized residuals, which indicate variables contributing disproportionately to the observed association (Tables S7 and S8). In the positive metagenome network, fatty acyls exhibited the largest contribution (χ² standardized residual = ±10.4), with temperate viruses displaying a greater-than-expected number of associations with this compound class (Fig. 5). In contrast, in the positive virome network, carboximidic acids and derivatives were the most strongly associated with temperate viruses (χ² residuals standardized residual = ±12.7).

**Figure 5.**
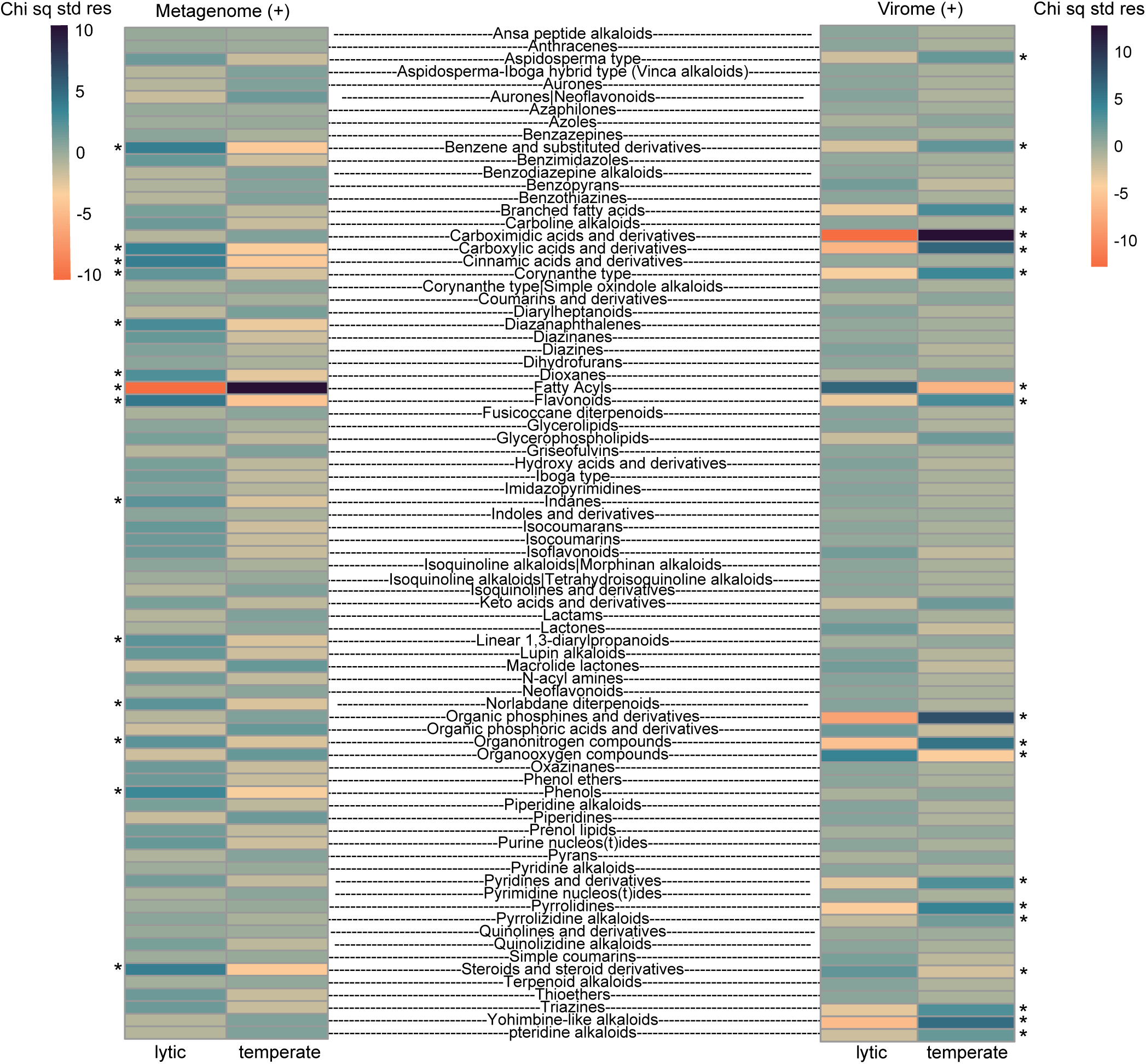
Associations between metabolites and viral replication strategy. The heatmap displays standardized residuals from a Chi-squared test with Monte Carlo simulations (50,000 iterations) for the positive metagenome virus-metabolite network (left) and the positive virome virus-metabolite network (right). Both networks exhibit significant differences in compound associations (χ² = 312.13, p < 0.0001 for the metagenome network; χ² = 588.27, p < 0.0001 for the virome network). Asterisks denote residuals > 2, which indicate compounds that contribute strongly to differences between temperate and lytic classes. The figure contains only shared compound classes across both networks.

The pattern observed for fatty acyls in the metagenome-derived network was reversed in the virome-derived network. By comparing high-ranking residuals (χ² residuals > 2) between the two networks, we found that most shared compound classes exhibited inverted associations. Specifically, benzene and substituted derivatives, carboxylic acids and derivatives, corynanthe-type alkaloids, flavonoids, and organonitrogen compounds were significantly associated with lytic viruses in the metagenome network but with temperate viruses in the virome network. Only one out of the seven compound classes, steroids and steroid derivatives, was significantly associated with the same viral lifestyle in both networks, displaying consistent patterns, though its effect size was relatively small in the virome network compared to other significant associations (χ² standardized residual = 2.32).

## Discussion

Here we show a significant relationship between the composition of the DOM pool and viral replication strategies on coral reefs. While the total DOM concentration was not significantly correlated with viral abundance, the network analyses show that the viral community displays significant associations with reef metabolites. The predominance of positive associations between viruses and metabolites (over 95% of the significant associations in both metagenome and virome networks) suggests that viral activity may either exert a top-down control on DOM and directly contribute to the production and transformation of metabolites or that a bottom-up effect takes place where metabolites indirectly select for different viral assemblages by affecting host assemblages. Cell-associated phages showed more frequent correlations with compounds of lower NOSC values (Fig. 2b), with temperate phages in particular associated with more reduced, energy-rich metabolites (Fig. 3c). This observation supports the notion that lysogeny may be selected for in an environment with an abundant supply of high-energy molecules. This is consistent with observations of low virus-to-microbe ratios on microbialized reefs with a supply of highly labile, algae-derived DOM (8, 21, 25).

### Top-down control: potential for viruses affecting the reef exometabolome composition

Phages may influence DOM composition through bacterial lysis (56, 57). However, tropical marine bacteria consist of approximately 12.4 fg of carbon per cell (58), and with cell densities of roughly 10^6^ cells per mL, contribute to about 1.03 µM of DOC. Considering that reefs in this study displayed on average 86.5 µM of DOC (SD =5.7), lysis of the entire bacterial community would still only make up 1.19% of the total DOC fraction. Coral reefs’ largest fractions of DOM are derived from macroalgae exudates and coral mucus (8, 12, 59). Therefore, it is unlikely that bacterial lysis would lead to observable changes in DOM.

Alternatively, viruses may influence DOM composition by enhancing or decreasing the efficiency of hosts metabolizing substrates, shifting their NOSC values. Phages encode auxiliary metabolic genes (AMGs) that augment host metabolism (30, 60–64). For instance, cyanophages can carry photosynthesis genes *psbA* and *psbD* to sustain energy production during infection, ensuring sufficient ATP generation for viral replication (61). Similarly, phage infection of *Pseudomonas aeruginosa* has enhanced pyrimidine and nucleotide sugar metabolism, suggesting metabolic reprogramming that supports both viral replication and host energy production (65).

Because bacteria are estimated to consume around 1.7-28.5% of DOC released by benthic primary producers (12, 66), it is possible that changes induced through viral infection or transduction may affect the nutrient uptake of bacteria and hence affect the DOM composition left behind. However, further experiments are required to test the prominence of these changes in bacterial consumption and whether they scale up to affect the composition of the total DOM pool.

### Evidence for compounds affecting viral replication strategy

A more likely scenario for the relationship between phages and metabolites is that the presence of energy-rich and highly reduced compounds may selectively favor specific bacterial taxa, which in turn affect the viral community. Low-NOSC compounds require higher oxidative power for degradation (67), and when energetic supply exceeds oxidative capacity, net ATP yield can decrease, leading to dissipation pathways and overflow metabolism (68, 69). The concept of increased electron donor to acceptor ratio (e^-^DAR) on coral reefs has been hypothesized to create conditions favorable for lysogeny (8, 12, 70). A higher e^-^DAR would result in ATP depletion coupled to an excess of NADPH, which promotes the accumulation of lytic phage repressors in model phage and bacteria (70). This may facilitate both the integration of new infections and the maintenance of existing prophages (68). Such a mechanism may explain the relationship between temperate viruses and more energy-rich compounds (Fig. 3d).

This mechanism is also consistent with the observation that the six most interconnected viruses within the metagenome-associated network were (Fig. 3c), despite the vast majority (> 90%) of the network consisting of lytic viruses. Notably, five of these viruses shared the same predicted host species, *Sphingobium yanoikuyae*, a heterotrophic bacterium frequently found in coral reef environments (28, 71, 72). *Sphingobium* species are well known for their ability to degrade environmental pollutants, including complex hydrocarbons, making them key candidates for bioremediation (73–75). For instance, the *Sphingobium yanoikuyae* B1 strain has been shown to break down polycyclic aromatic hydrocarbons (PAHs) in contaminated soils (73). Aromatic degradation requires substantial molecular oxygen and energy input (76) and may lead to ATP depletion and oxygen limitation, potentially creating favorable conditions for phage integration. In a previous proximity-ligation study, five viruses infecting *Sphingobium yanoikuyae* displayed a range of ecological strategies, from latent to highly productive, as inferred from their relationships with host abundance (26). This variability suggests that these phages may transition between different infection strategies, with a predominant tendency toward long-term lysogeny, which could be explained by their host’s metabolism.

Chi-squared analyses revealed a significant association between viral lifestyle and compound class across both networks, with the compounds contributing most strongly to this pattern exhibiting opposite associations with lytic and temperate viruses in the metagenome-versus virome-derived datasets (Fig. 5). This may further reflect evidence for compounds affecting viral replication strategies. A temperate virus must undergo lysis to be abundant in seawater, whereas its presence in the cell-associated fraction suggests lysogenic infection. Therefore, a highly abundant temperate phage in seawater likely indicates a lytic lifestyle, while its prevalence in the cell-associated fraction points to lysogeny. The stronger relationship observed between energy-rich compounds and cell-associated viral abundances (Fig. 2b), but not viromes (Fig. 2c), may reflect that metagenome-derived viruses are more tightly linked to host abundances (26), including integrated prophages. This distinction helps explain the observed inverse relationship between compounds in these fractions. Therefore, both independent networks highlight similar trends between lytic and lysogenic cycles.

### Implications for coral reef biogeochemistry and functioning

Our findings have implications for coral health within the DDAM framework by potentially supporting the feedback loop. Fleshy algae exudates are highly labile and energy-rich (47), driving shifts in microbial community composition and increasing microbial abundance (15–17, 77). The increased labile DOC supply has been hypothesized to increase lysogeny through the e^-^DAR mechanism described above (68). Although we did not identify the source of each compound, our results support this hypothesis by showing that high-energy compounds were associated with viral signals indicative of lysogeny (Fig. 2b, Fig. 3c,d). Compounds with high Gibbs energy values have also been associated with the enhanced growth of opportunistic coral pathogens (78), further reinforcing the potential for coral disease. In Pacific coral reefs, a low virus-to-microbe ratio, a proxy for lysogeny, has been associated with low coral and increased algae cover (7). Combined with previous observations, we suggest that viruses may reinforce the DDAM feedback loop by shifting viral lifestyles toward lysogeny, fostering the growth of copiotrophic and potentially pathogenic bacteria. In conclusion, our findings show a significant relationship between temperate phages and energy-rich DOM. This may exacerbate the DDAM feedback by allowing the persistence of virulent microbial communities through lysogeny.

## Acknowledgments

We thank Prof. Dr. Mark Vermeij, Dr. Jason Bear, and Lars ter Horst for their field support at the CARMABI research station in Curaçao. We would like to thank Denise Dorhout, Monique Verweij and the Lipid Lab at NIOZ for their expertise and support during exometabolomic data acquisition.

## Author’s contributions

CBS and NSV conceptualized the work. NSV, CSB, AFH, WB, and YS collected samples. NSV, CSB, YS, WB, and LS processed samples. NSV and LS performed analyses. NSV wrote the first manuscript version, and all authors contributed significantly to the manuscript.

## Supplementary Tables

**Table S1:** List of geographic locations and sample types collected. **Table S2** Viral genomes identified as temperate. **Table S3** Significant Pearson correlations adjusted for multiple comparisons (p < 0.01) between free viruses and metabolites. **Table S4** Significant Pearson correlations adjusted for multiple comparisons (p < 0.01) between metagenome viruses and metabolites. **Table S5** Abundances of superclass analogs detected by the np classifier. **Table S6** IPHop results of the 10 viral genomes most frequently associated with metabolites in the cell-associated and free virus networks. **Table S7** Counts and residuals from Chi-squared test with Monte Carlo correction (p-value = 2e-05), testing differences between viral lifestyles and compound associations for cell-associated viruses. **Table S8** Counts and residuals from Chi-squared test with Monte Carlo correction (p-value = 9.999e-05), testing differences between viral lifestyles and compound associations for free viruses.

